# Statistical determinants of visuomotor adaption along different dimensions during naturalistic 3D reaches

**DOI:** 10.1101/2021.02.16.431440

**Authors:** E. Ferrea, J. Franke, P. Morel, A. Gail

## Abstract

Neurorehabilitation in patients suffering from motor deficits relies on relearning or re-adapting motor skills. Yet our understanding of motor learning is based mostly on results from one or two-dimensional experimental paradigms with highly confined movements. Since everyday movements are conducted in three-dimensional space, it is important to further our understanding about the effect that gravitational forces or perceptual anisotropy might or might not have on motor learning along all different dimensions relative to the body. Here we test how well existing concepts of motor learning generalize to movements in 3D. We ask how a subject’s variability in movement planning and sensory perception influences motor adaptation along three different body axes. To extract variability and relate it to adaptation rate, we employed a novel hierarchical two-state space model using Bayesian modeling via Hamiltonian Monte Carlo procedures. Our results show that differences in adaptation rate occur between the coronal, sagittal and horizontal planes and can be explained by the Kalman gain, i.e., a statistically optimal solution integrating planning and sensory information weighted by the inverse of their variability. This indicates that optimal integration theory for error correction holds for 3D movements and explains adaptation rate variation between movements in different planes.

## Introduction

Studies of motor control and motor adaptation previously have been performed mostly in one or two-dimensional settings^1–7^. It is unclear, how well these findings generalize to movements in 3D, since 3D settings apply much less physical constraints on body pose and movements^8,9^. Unless movements are constrained in a horizontal plane by a table or supported by an exoskeleton, gravity induces a force anisotropy; unless movements are conducted in a frontoparallel plane, visual depth induces a perceptual anisotropy. It is unknown, if and how differences in the way subjects plan movements and in the way they perceive the environment and their own movement along different dimensions translate into motor adaptation anisotropy. Here we directly compared adaptation in different dimensions in the context of naturalistic movements in 3D.

Perceptual as well as other factors might contribute to adaptation anisotropies. Studies in 3D virtual reality (VR) found that adaptation along a vertical axis entails a reduced learning rate relative to the other two axes when a simultaneous triaxial perturbation was applied^10^. The authors proposed that their results are a consequence of the higher weight given to proprioception when adjusting movements along the vertical axis compared to sagittal or horizontal axis, considering/assuming that motor learning based on proprioceptive feedback entails a lower learning rate than based on visual feedback. We will refer the variability attributed to visual or proprioceptive feedback and affecting the estimate of the effector (hand) position as “measurement variability”^11^. However, not only measurement variability matters. According to optimal feedback control theory (OFC), the nervous system optimally (with respect to a specified goal) combines sensory feedback (e.g. visual and proprioceptive) information with a forward prediction of the body’s state to control movements and to correct errors^12–15^. During motor adaptation, this forward model is updated to reduce a sensory prediction error arising from a mismatch between its prediction and the sensory feedback. Any variability in this closed-loop process thus should affect the observed rate of adaptation. This includes variability in movement planning^11,16–21^ which might result from the stochastic nature of neuronal processes in sensorimotor transformations, referred to as “planning variability”, and which directly affects the reliability of the forward model prediction^22–26^. Here, we test the prediction of optimal integration theory that planning and measurement variability between different dimensions relative to the body determine the corresponding motor adaptation rates^16–19,27^.

According to optimal integration theory, a statistically optimal solution for the update of the forward model during adaptation is the Kalman filter^28^, which is updated from the error proportionally to the inverse of measurement variability and planning variability^11,29,30^. Higher variability in the forward model (high planning variability) would increase the Kalman gain and allow faster corrections of the experienced error, whereas a higher variability in the sensory feedback (high measurement variability) would decrease the gain and therefore lead to slower adaptation. Previous studies found that human behavior approaches that of an optimal learner^11,21,31^ showing that adaptation rate positively correlates with the Kalman gain calculated from the planning and measurement variabilities. Also, individual differences between subjects in the rate of motor adaptation can be largely explained by individual differences in the Kalman gain^20^, a prediction which we here test in the context of 3D movements. To study adaptation in naturalistic 3D movements, we leverage existing differences in planning and measurement variability in the different planes of space (coronal, sagittal, horizontal) instead of experimentally inducing variability. From previous work, we expect that the higher variability associated with depth perception reduces the rate of adaptation ^14,32^. At the same time, planning variability might be elevated along the vertical axis given the stronger proprioceptive rather than visual component ^33^ due to the need of compensating gravitational forces ^34–36^.

Current experiments and models propose that sensorimotor adaptation is supported by two learning processes, which act on different timescales, one fast adapting and fast forgetting and one slow adapting and slow forgetting^7,37^. To estimate adaptation rates separately for each learning process and link them to Kalman gain, we will fit both fast- and slow-state processes during sensorimotor adaptation. Compared to one-state models^22,23,26^, two-state models explain motor adaptation as a combination of explicit and implicit learning mechanisms, where explicit learning allows fast adaptation and implicit learning changes performance more slowly^4,6,38^. Two-state models also better explain phenomena like savings, i.e. improved learning due to repeated exposure^39–41^, and anterograde interference, i.e. the negative influence of past learning on future learning^2,42^, that was found to be partially correlated with explicit learning^5^. Moreover, neurophysiological evidence of two-state learning dynamics was found in humans and monkeys^43,44^. In a VR context, explicit learning was found to have a stronger influence on the final adaptation level than in non-VR adaptation paradigm^8^. Given that two-state dynamics were found to better model adaptive behaviours of visuomotor rotation paradigms, especially in VR settings, our method will be focused on a two-state space model but the results will be also compared with a one-state model.

By considering the stochastic nature of error-based learning, we can independently estimate the measurement and planning uncertainty from the empirical movement data during a 3D adaptation task using stochastic modelling. Stochastic modeling has been shown to better identify the time course of the hidden states^45^. Specifically, we use Bayesian inference via Hamiltonian Markov chain Monte Carlo sampling (referred to as Hamiltonian Monte Carlo, HMC) to implement a novel multilevel hierarchical version of the two-state space model fitting. Hierarchical modelling generally allows greater regularization of the subjects’ parameters and simultaneous estimates of the parameter distributions ultimately resulting in less overfitting of the data^46^. We evaluated the hierarchical model and compared it to another state-of-the-art model performing Bayesian inference via an expectation-maximization (EM) algorithm^45^ without a hierarchical structure. We show that HMC allows for a better extraction of data on the two underlying motor learning states and the planning and measurement variabilities. Using hierarchical HMC models, we confirm that two-state models better explain our experimental three-dimensional data than single-state models. We, therefore, use the two-state hierarchical HMC model to test whether differences in adaptation dynamics can be found between movements and perturbations of different directions relative to the subject’s own body and whether these differences comply with optimal integration theory.

## Materials and Methods

### Subjects

Data were collected from 26 healthy subjects (age range 20-32, 10 females, 16 males). The study was performed under institutional guidelines for experiments with humans, adhered to the principles of the Declaration of Helsinki, and was approved by the ethics committee of the Georg-Elias-Mueller-Institute for Psychology at the University of Goettingen. All the subjects were right-handed, had normal or corrected-to-normal vision and received financial compensation for their participation. Subjects gave written consent to participate and received detailed written instructions for the task. They were asked to paraphrase the instructions in their own words to make sure they understood the task. The experiment was carried out in one single appointment that lasted between 2.5 and 3 hours, depending on the subjects’ freely chosen breaks among blocks (see later).

### Virtual Reality Setup

Subjects sat in a darkened room at typical room temperature with no distracting noises. The 26 experiments were done individually at different times of the day and on different days. Subjects performed the visuomotor task in a three-dimensional (3D) virtual reality with Wheatstone stereoscopic view. Visual displays on two monitors were controlled by a customized C++ software similar to what was described in previous studies ^47,48^. The images of the two monitors (one for each eye, BenQ XL2720T, 27-inch diagonal, 1920×1080, 60 Hz refresh rate, BenQ, Taepei, Taiwan) were reflected by two semi-transparent mirrors. (75 x 75 mm, 70/30 reflection/transmission plate beamsplitter, covered from the back to block transmission; stock #46-643, Edmund Optics Inc., Barrington, New Jersey, United States) that were angled 45° relative to the two monitors on each side of the subject (Fig. 1A).

**Figure 1.**
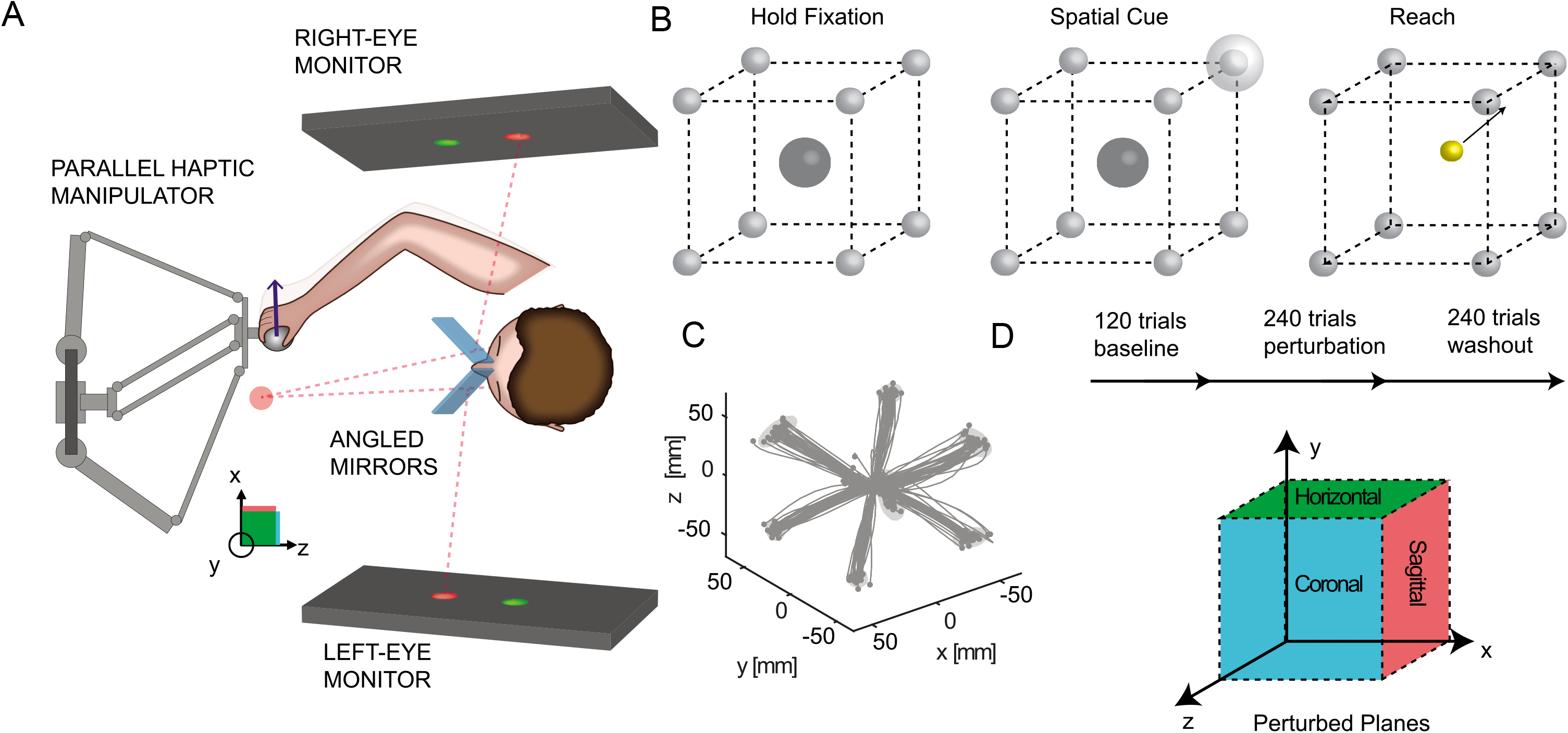
Virtual reality setup with robotic manipulator for testing visuomotor rotation adaptation of naturalistic reaches in different planes relative to the body. A) Schematic representation of the 3D virtual reality setup. The two angled mirrors reflect the image from the two screens to the two eyes independently providing stereoscopic vision. The mirrors are semitransparent to allow calibration of the virtual reality space with the real hand position. During the experiments, transparency is blocked, so that the subjects do not see the robotic manipulator and their hand, but only the co-registered cursor. B) Spatial and temporal structure of the 3D reaching task. Following a brief holding period in the center of a virtual cube, the subject has to reach to one of eight targets arranged at the corners of the cube. For ergonomic reasons, the cube is orientated relatively to the slightly downward-tilted workspace and head of the subject, not according to true horizontal. C) Example baseline trajectories of a single subject. D) Time course of the perturbation experiment. Following a baseline phase of 120 trials, a 30-degree visuomotor perturbation is applied for 240 trials, followed by a washout of 240 additional trials. In three separate blocks for each subject, the perturbation is applied in the sagittal, coronal and horizontal plane, respectively. Note that all reaches are conducted along a diagonal direction, i.e., each movement is affected by either of the perturbations.

The two screens render a 3D view of virtual objects on a dark background. To accommodate test subject ergonomics, we tilted the plane of view (mirrors and monitors) downward from horizontal by 30° towards the frontal direction (virtual horizontal). This ensured that arm movements in the physical space could be carried out in an ergonomic posture in front of the body while the virtual representation of the hand (the cursor) was aligned with the actual hand position. The virtual cube that defined the subjects’ workspace was centered on and aligned with the axis of gaze. For the rest of the paper, we describe orientation relative to the axes of this cube, i.e. relative to the virtual, not the physical horizontal and vertical.

To optimize the 3D perception of the virtual workspace for each subject, the inter-pupillary distance was measured (range: 51 to 68 mm, mean = 59.48 mm) and the 3D projection was modified in the custom-made software accordingly. This has been shown to benefit depth perception, reduce image fusion problems and reduce fatigue in participants^49^.

Subjects performed controlled physical arm movements to control cursor movements in the 3D environment. For this, they moved a robotic manipulator (delta.3, Force Dimension, Nyon, Switzerland) with their dominant hand to place the 3D cursor position into visual target spheres. The manipulator captured the hand position at a sampling frequency of 2 kHz with a 0.02 mm resolution. Subjects could neither see their hand nor the robot handle. Instead, a cursor (yellow sphere, 6 mm diameter) indicated the virtual representation of the robot handle and therefore the hand position (Fig. 1B).

### Behavioral task

Subjects performed a 3D center-out reaching task, i.e. started their reaches from a central fixation position (indicated by sphere, semitransparent grey, 25 mm diameter) to one of eight potential target positions (grey sphere, 8 mm diameter) arranged at the corners of a 10 cm-sided cube centered on the fixation position (Fig. 1B–C). Each subject completed three blocks with breaks in between. One block consisted of 600 successful trials: 120 baseline trials, then 240 perturbed trials, and 240 washout trials again without a perturbation (Fig. 1D). Each block lasted between 20 to 25 minutes, depending on the performance of the subject. Breaks between blocks were adjusted according to the subject’s needs and usually lasted between 10 to 15 minutes for compensating potential fatigue/savings/anterograde interference effects (see later). Before the actual experiment, a 10-minute training session was run to make subjects comfortable with the three-dimensional environment and to ensure a correct understanding and execution of the task.

At the beginning of each trial, subjects moved the cursor (co-localized with the physical hand position) into the central fixation sphere and held the cursor there for 200 ms (“hold fixation”; Fig. 1B). After this holding period, one of the eight potential targets was briefly highlighted by a spatial cue (semitransparent grey sphere of 25 mm diameter, centered on the target). The corners of the cube, and hence also the target, remained visible after the cue stimulus disappeared to allow precise localization of the target in the 3D space. The size of the spatial cue corresponded to the eligible area in which the movement had to end to be considered successful. The spatial cue remained visible for 300 ms and then disappeared together with the fixation sphere. This disappearance served as a GO signal for the subjects. Subjects then had 500 ms time to leave the fixation sphere and begin their hand movement towards the target. For successful completion of the trial, it was sufficient that the cursor completely entered the target sphere, without the need to stop inside the eligible area. Success or failure was indicated by sound signals: high-pitch for success, low-pitch for failure. A trial was aborted if subjects had not left the starting sphere 500 ms after the GO signal, if the movement was stopped (speed below 3cm/s) at any point outside the target area or lasted longer than 10 seconds. After each trial, subjects could immediately start a new trial by moving the cursor back to the starting position.

Within each block, we consistently applied the same 2D visuomotor perturbation to the 3D position of the cursor during all perturbation trials. Each perturbation affected the cursor position either within the xy-, the yz- or the xz-plane, respectively. This means, for every perturbation, one out of the three dimensions was left unperturbed, corresponding to the axis of rotation for the perturbation. In the perturbed trials, the workspace was rotated by 30 degrees counterclockwise around either the x (rotation in the sagittal plane), y (rotation in the horizontal plane) or z (rotation in the coronal plane) axes in each of the three blocks (Fig. 1D). The rotation was applied in each block to one of the three planes. As proof of concept, we verified that the projections of the trajectories on the perturbed planes look similar (Fig. 2A). The order of the blocks (perturbation planes) were controlled over the experiment using stratified randomization between subjects, to prevent potential effects of fatigue, interference or savings between blocks.

**Figure 2.**
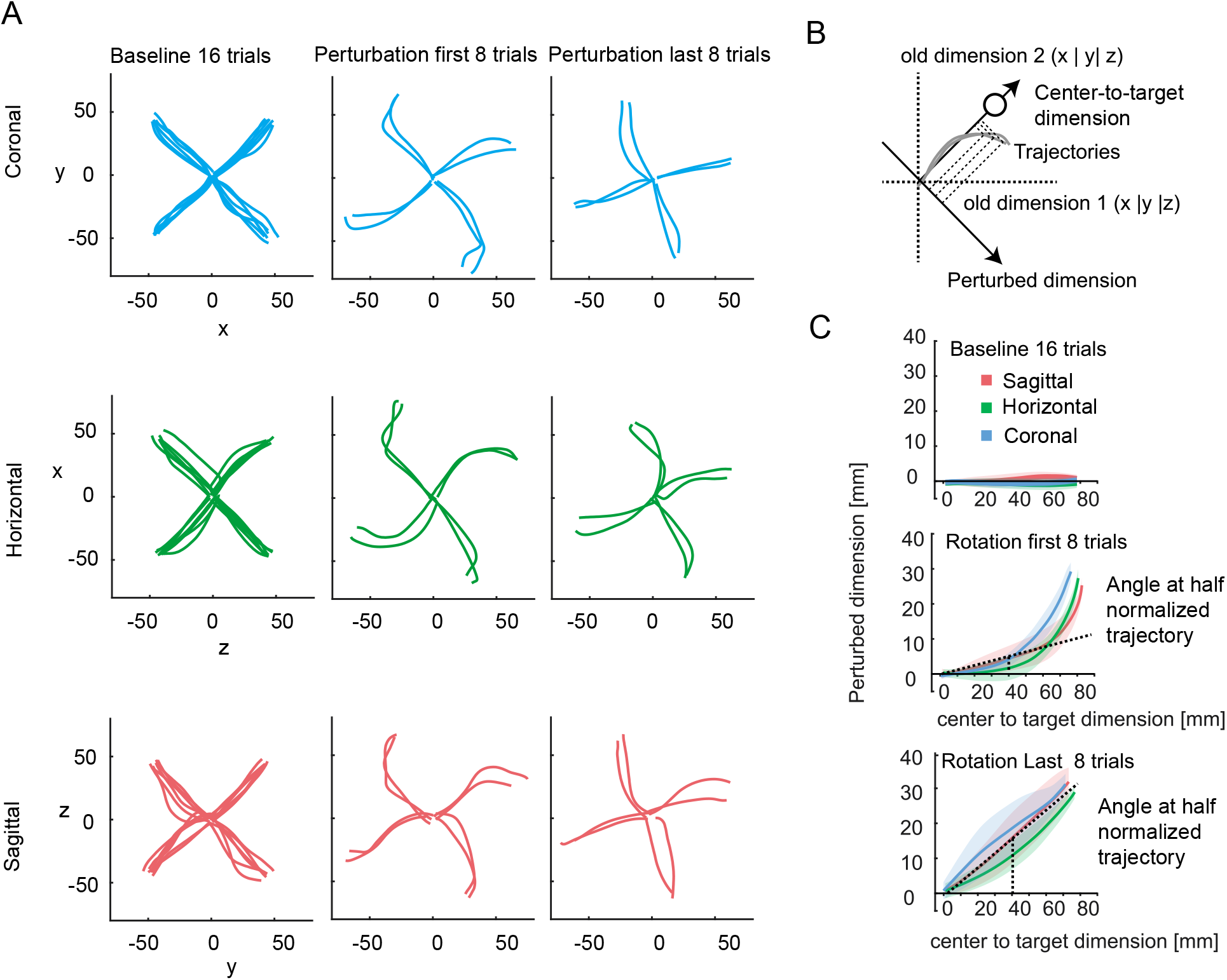
Quantification of adaptation level. A) Example trajectories from one subject. Each row/color of the graph correspond to different planes of rotation. The trajectories are projected onto the plane which is perpendicular to the axis of visuomotor rotation for representation and analysis of the data in each of the three conditions (coronal, horizontal, sagittal). The trajectories show similarities across the three planes. B) Within each plane, the trajectories are quantified along the two task relevant dimensions, the center-to-target dimension and the dimension orthogonal to it in the same plane, which we refer to as “perturbed” dimension. To compare data across tasks, each trajectory was resampled and rotated into this reference frame. At half of the normalized trajectory, the adaptation angle was calculated and used for model fitting. C) Examples of transformed trajectories from one subject during baseline, early adaptation and late adaptation.

In the three blocks, the subjects conducted the same movements in terms of starting and end positions. We chose the corners of a cube as targets since any movement from the center of the cube to a corner always is a diagonal movement that is composed of equally large x-, y- and z-components. Therefore, the same target movement can be subjected to perturbations in each of the three planes parallel to the surfaces of the cube. This allowed us to compare perturbations in different body-centered dimensions without changing movement start and end positions, thereby avoiding potentially confounding effects of posture.

Within each block of trials, the order of the targets was pseudo-randomized such that all eight targets were presented within each set of eight successful trials. These 8-trial sets counted as epochs and the model fitting analysis was carried out on epochs rather than on single trials.

### Data preprocessing

Data were stored for offline analysis performed in MATLAB^®^ R2018a (MathWorks, Inc., Natick, Massachusetts, United States). Data plots were generated with gramm, a MATLAB plotting library^50^. Trajectories were aligned in time to the start of the movement, defined as a speed increase above 3 cm/s, to when the hand reached the target, and resampled in 50 time steps, so that each trajectory was normalized in duration.

To calculate the spatial deviation of the hand position for each perturbed plane independent of target direction, the trajectory of each trial was projected on the axis orthogonal to the center-to-target direction and laying in the plane of the applied perturbation (Fig. 2B, perturbed dimension). To remove biases that would result from curved trajectories independent of any perturbation, we computed the average trajectory during baseline trials for each subject and block and subtracted these mean trajectories from each trajectory in the according block. Figure 2C (middle and bottom panel) shows, for one example subject, the average of the projected trajectories at baseline, during the first eight perturbation trials (early adaptation) and the last eight perturbation trials (late adaptation) for all three perturbed planes. While baseline-corrected trajectories are straight during baseline (by construction), the early adaptation phase is characterized by curved trajectories with online movement corrections to reduce the target error induced by the perturbed visual feedback. Later during adaptation trajectories on average were straight again, i.e. the subject aimed at a direction suited to compensate the effect of perturbed feedback from beginning of the movement. To quantify changes of the initial aiming direction over the course of adaptation, we measure the deviation of the trajectory along the perturbed dimension at the moment when half the distance along the center-to-target dimension is travelled (Fig. 2C) and quantify it as angle.

### Fitting model and procedure

Existing Bayesian tools for fitting adaptation models^20,45^ can only estimate parameters from single experimental runs (changes over repeated trials in one task condition and one subject) despite the often hierarchical nature of the experiment (repeated trials interleaved with multiple conditions, multiple subjects).

To get more reliable estimates of subject-level and population-level learning parameters, we developed a hierarchical-model version of single-state and two-state models of adaptation. Hierarchical models for Bayesian inference allow to regularize the single subject fitted values by introducing a hyper-parameter at the population level. For this, we used the probabilistic programming language Stan^51^, through its MATLAB interface. Stan allowed us to perform Bayesian inference with its implementation of a Hamiltonian Markov chain Monte Carlo method using a NUTS sampler algorithm^52^ for fitting the model parameters to the empirical data.

The underlying trial-by-trial adaptation model for the movement angle corrections is the following. For the trial *n* in subject *j* and perturbation plane *p*, the measured movement angle *y*_*jp*_(*n*) is:

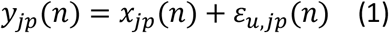

With *x*_*jp*_ (*n*) being the hidden internal state (planning angle) and *ε*_*u,jp*_ (*n*) the measurement noise of the movement due to limited sensory accuracy of the agent, described by a normal distribution with zero mean: *ε*_*u,jp*_ (*n*)~*N*(0, *σ*_*u,jp*_). We refer to this term as measurement noise since it reflects the uncertainty of estimating the position of the hand based on the sensory feedback with zero mean and standard deviation *σ*_*u,jp*_ from which we calculated the measurement variability by squaring the term *σ*_*u,jp*_ 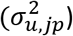. We use this term to differentiate it from the uncertainty arising at the central brain level (planning noise; see below). In our experimental paradigm/model, the measurement noise is also influenced and cannot be disentangled from the execution noise that is added to the movement during conduction at the “periphery”, i.e., it includes any movement variability that is accessible to the central nervous representations only via sensory feedback.

The internal state *x*_*jp*_(*n* + 1) in trial *n* + 1 depends on the state in the previous trial *n*, with retention **A**, and the error ***e*** in the previous trial, with learning rate **B**. The error *e* is expressed as **e**_jp_(n) = **p** − **y**_jp_(**n**), where p is equal to thirty degrees during perturbed trials and zero degrees otherwise. The state update is affected by a planning noise (or state update error) *ε*_*x*_(*n*), which gives us the following:

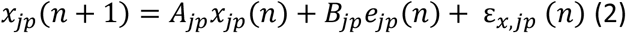

In the single-state model, *A*_*jp*_and *B*_*jp*_ are positive scalars, and *ε*_*x,jp*_(*n*)~*N*(0, *σ*_*x,jp*_). *σ*_*x,jp*_ is the standard deviation of the internal state error and allows calculating the planning noise by squaring the term *σ*_*x,jp*_ 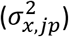. For the two-state model:

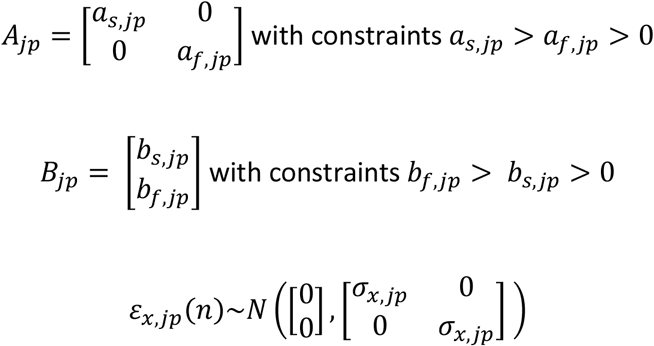

In our model, all subject and plane-level parameters: *σ*_*u,jp*_, *a*_*s,jp*_, *a*_*f,jp*_, *b*_*s,jp*_, *b*_*f,jp*_, *σ*_*x,jp*_ were drawn from population-level normal distributions, e.g. for *a*_*s,jp*_ we define 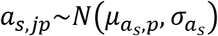. Here, 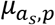 and 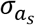 are population-level hyper-parameters, corresponding to the population (between-subject) average of the slow retention for perturbation plane *p* and the population standard deviation of the slow retention.

To keep the model simple, we did not include common per-subject offsets in the average hyper-parameters across planes; in other words, the model we used did not take into account the fact that the same subjects participated in experimental runs with different perturbed planes. A variation of the model which permitted per-subject offsets yielded equivalent results (data not shown).

Bayesian inference requires priors for the hyper-parameters, which we chose wide to not constrain the model. Retention average hyper-parameter (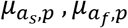 in the two-state model and *μ*_*A,p*_ in the single-state model) priors were ~*N*(1,9), learning rate average (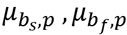 and *μ*_*B,p*_) priors were ~*N*(0,9), and their population standard deviation hyper-parameter (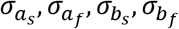 and *σ*_*A*_, *σ*_*B*_) priors were half-Cauchy distributions (location = 0, scale = 5)^53^. Priors of the population average and standard deviation hyper-parameters of the measurement and planning noise (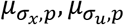 and 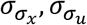) were half-Cauchy distributions (location = 0, scale = 15). The conclusions from the model fitting do not critically depend on the exact choice of any of these parameters.

### Steady-state Kalman gain

From the variances of the noise processes, we calculated the steady-state Kalman gain, as derived by Burge and colleagues^11^. In short, by combining measurement and state-update equations of the Kalman filter and assuming that a steady-state is reached, we can introduce the term 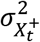 which consist on the variance of the best estimates given the two noisy measurements (see Burge et al., 2008, for full formula derivation):

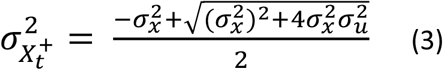

The Kalman gain (K), weighting the contribution of planning and measurement variabilities in error correction, is calculated as follows and indicates the amount of correction that should be attributed to the error:

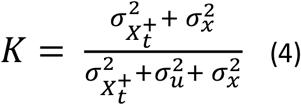

### Validation of fitting procedure

To validate our hierarchical Bayesian fitting algorithm, we compared it to a non-hierarchical algorithm by applying both to surrogate data sets. For this, we generated N = 100 artificial experimental runs using an underlying two-state time course with the same number of baseline, perturbation, and washout trials as in our experiment. For each simulation, each parameter was drawn from a random normal distribution approaching the values we observed in our data: as~N(0.93,0.03), af~N(0.55,0.2), bs~N(0.06,0.04), bf~N(0.18,0.08), planning variability~N(1.5,1), measurement variability~N(6,3). We then compared the accuracy of parameter estimations from different fitting methods, namely an EM algorithm ^45^ and a non-hierarchical and a hierarchical HMC algorithm.

To quantify whether our two-state hierarchical algorithm introduces spurious correlations among the learning rates and the Kalman gain, we applied it to two versions of the surrogate datasets: One dataset was built with 200 completely uncorrelated samples, whereas in a second data set we imposed bs equal to the 0.3 times the Kalman gain calculated according to (4).

## Results

The subjects performed the task (three conditions: baseline, perturbation and washout) with a high success rate in all perturbed planes (number of correct trials/number of total trials > 99%). We used a generalized linear mixed effect model (GLMM) with a binomial distribution to test the effect of the perturbation in the different planes (fixed effect) on the success rate, with the subjects as random effect. With the sagittal plane as intercept, there was no significant effect on the slope for the horizontal plane (tstat = 1.5477, p = 0.1217) and no significant effect for the slope of the coronal plane (tstat = 1.3976, p = 0.16223). This means that we did not observe differences in performance among the three perturbation planes.

The subjects showed stereotypical adaptation time courses (Figure 3A and supplementary figures 1–3) during and after introduction of the perturbation with typical error correction profiles that compensate for the applied rotation. To test differences among the three planes regarding the level of adaptation reached by the subjects (Fig. 3A), a one-way between-subjects ANOVA was conducted to compare the effects of the perturbed plane over the final adaptation level. The final adaptation level was calculated as the average of the adaptation time course over the last 4 epochs of the perturbation period (Fig. 3B). We found that the subjects adapt differently to the perturbations depending on the plane of rotation (F(2,69) = 30.77, p = 2.8 E-10). The final level of adaptation was significantly higher for perturbations in the coronal plane compared to the sagittal plane (coronal-sagittal comparison, p = 7.6E-05, post-hoc multi-comparison test corrected with Tukey’s honestly significant difference) and horizontal plane (coronal-horizontal comparison, p = 1.0 E-08) (Fig. 3B). Moreover, perturbations in the sagittal plane provoke stronger adaptation than perturbations in the horizontal plane (sagittal-horizontal comparison, p = 0.004) (Fig. 3B). To understand the underlying dynamic of the differences in adaptation, we modeled adaptation behavior in the three different visuomotor perturbation planes with a two-state model, which separately identified the slow and fast processes underlying adaptation. While our hierarchical Bayesian model fits data across conditions and subjects (see Methods), Figure 3C shows the fitted curves for one example subject performing three adaptation conditions.

**Figure 3.**
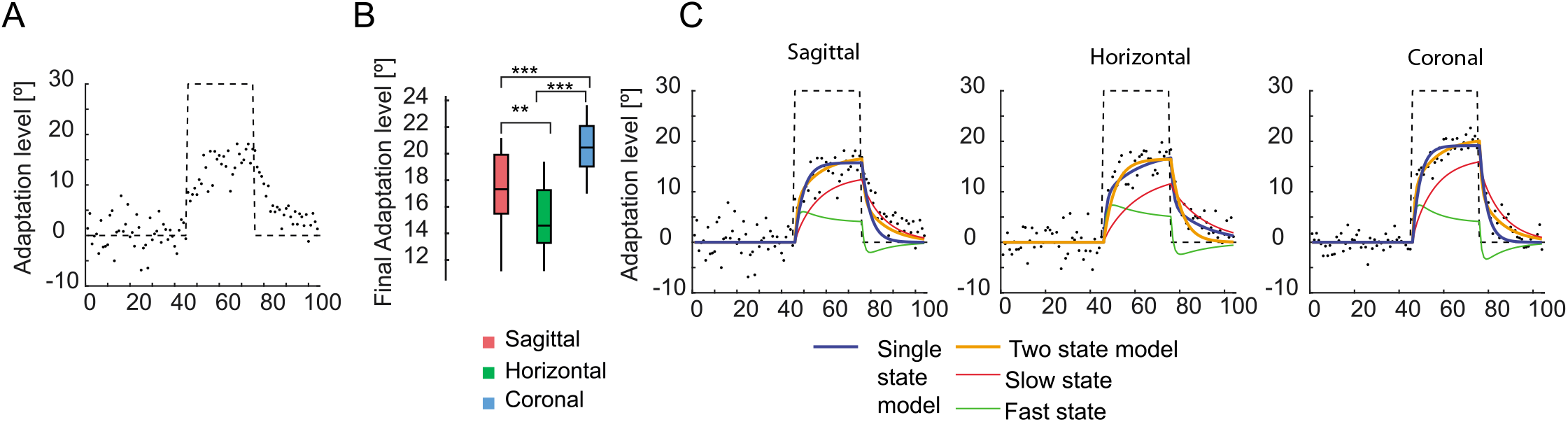
Visuomotor adaptation in three different planes relative to the body. A) Example of the time course of adaptation for one example subject with perturbation applied to the sagittal plane. B) Final adaptation level for each plane. C) Examples of data fitted with a one-state and a two-states learning model, respectively. Black points represent the subject’s data (average movement angle at half-trajectory within 8-trial epochs; 105 epochs represent 360 concatenated baseline trials, 240 adaptation trials and 240 washout trials). The black dashed line shows the perturbation profile.

Before comparing adaptation between the different planes, we evaluated whether in our three-dimensional task, a two-state model fits the data better than a single-state model. The single-state model shows larger root-mean-square residuals, i.e., larger difference between modeled and observed adaptation trajectory (paired t-test, p = 6.6e-16). To take into account the number of free parameters when comparing models, we additionally computed the widely applicable information criterion (WAIC), an approximation of cross-validation for Bayesian procedures^54^. The two-state model shows a lower WAIC compared to the single-state model (3.54E+04 and 3.58e+04, respectively), confirming the advantage on using a two-stage model.

To validate our hierarchical fitting algorithm, we tested whether model parameters are better reconstructed by our two-state Bayesian HMC hierarchical algorithm compared to a state-of-the-art EM algorithm^45^. We also compared the HMC hierarchical model with the corresponding non-hierarchical HMC model. For this, we generated multiple surrogate two-state adaptation datasets with randomly sampled parameters (see Methods). We then tested how well the three algorithms retrieve the parameters of the surrogate datasets. The hierarchical algorithm better reconstructs all parameters (error is smaller and correlation is higher) (Fig. 4). We therefore decided to use a two-state model and independently fit the parameters for each perturbation plane.

**Figure 4.**
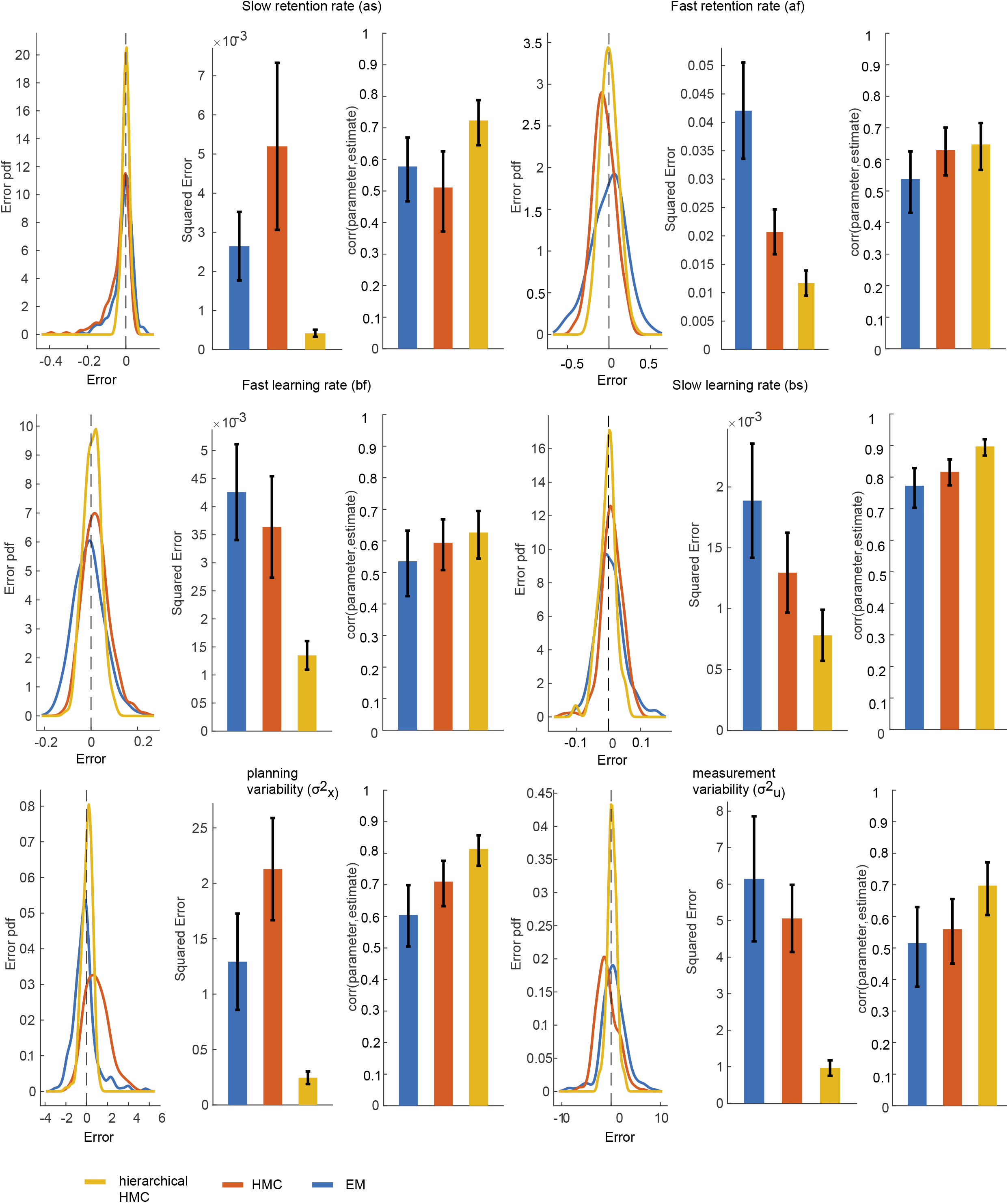
Comparison of hierarchical and non-hierarchical fitting algorithms. Parameter estimation quality for our HMC hierarchical fitting procedure (orange), an HMC non-hierarchical fitting procedure (red), and a EM fitting procedure (EM, blue; ^45^). For each random parameter setting of the surrogate two-state datasets, we represent the error in parameter reconstruction (estimated minus set parameter) for each fitting procedure, including the distributions of the errors, average and 95% confidence intervals of the squared error; and the correlation (average and 95% CI) between the estimated and set parameters.

Optimal integration (OI) theory predicts that the rate of adaptation correlates with the weighing between internal estimation and external sensory information for assessing the state of the system. A learner optimally integrating these two types of information approximates a Kalman filter that combines them depending on their respective variabilities. If planning (internal estimate) variability is high and measurement (sensory information) variability low during adaptation, the experienced error will be given credibility, and therefore faster learning will happen. In the opposite case, the learner tends to disregard the experienced error and corrections would happen slowly.

We fitted all learning parameters of the two-state model and the measurement and planning variabilities from our empirical data (Fig. 5A). To test if the above-mentioned mechanism of OI can explain the observed differences between the perturbed planes in our experiment, we also correlated planning and measurement variabilities with the slow learning rate bs for each experimental run in our dataset (Fig. 5B). Correlations between planning variability and the learning rate as well as between measurement variability and learning rate were low (respective R2 : 0.13 and 0.11), but with a trend consistent to the OI predictions, i.e., low measurement noise and high planning noise correlate with higher values of the slow learning rate. When combining the contribution of the two noise parameters into an estimate of a Kalman gain for each subject, the slow learning rate was explained much better (R2 between K and bs: 0.48), with a high Kalman gain being associated with high learning rate. Notably, this correlation between Kalman gain (K) and slow learning rate (bs) is determined by the different levels for K and bs between the perturbation planes. This means, the slow learning rate is mostly explained by the plane of the workspace in which the perturbation is applied (R2 = 0.94), while it does not correlate with the Kalman gain variation across the individual subjects within a perturbation condition. Last, the fast learning component shows a correlation with the Kalman gain among perturbed planes (R2 between K and bf: 0.34).

**Figure 5.**
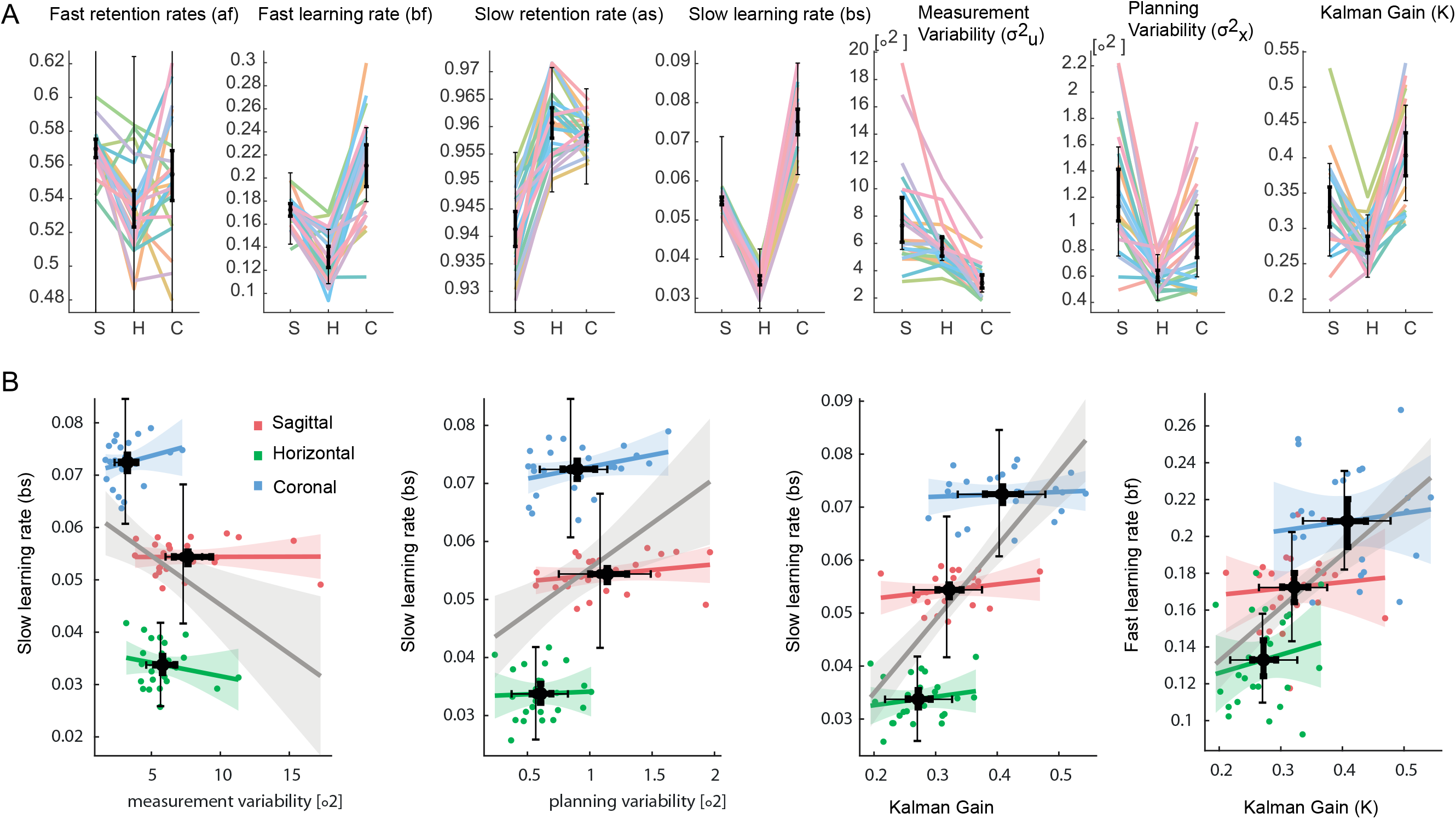
Comparison of two-state model parameters between the perturbation planes. A) Estimated fast and slow retention (*a*_*f,jp*_, *a*_*s,jp*_), learning rates (*b*_*f,jp*_, *b*_*s,jp*_), and measurement and planning variabilities (*σ*_*x,jp*_, *σ*_*u,jp*_) for the sagittal (S), horizontal (H) and coronal (C) perturbation planes. Each color represents one subjects. The error bars correspond to 95 % credible intervals (i.e., contain 95% of the posterior probability) for the population average hyper-parameters 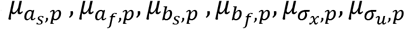. The steady-state Kalman gain K was computed from the posteriors of the measurement noise and planning noise. B) Relationships between the slow learning rate bs and variability related parameters: measurement noise variability, planning noise variability, and steady-state Kalman gain. Points correspond to individual subjects’ parameters in the different rotation planes (coded by color). The correspondingly colored lines represent linear fits based on the across-subject parameter values within each plane. The grey line corresponds to a linear fit based on subjects’ parameters values across all planes. The horizontal and vertical error bars correspond to 95 % credible intervals for the corresponding parameter population average hyper-parameters.

A long washout plus additional waiting time between conditions prevented savings or anterograde effects when switching between different planes of perturbation. To test for this, a forward stepwise linear regression was used to identify possible predictors of the learning rates (bf or bs independently) out of the order of execution (1st executed task, 2nd execution task or 3rd executed task) and perturbed axis. At each step, variables were added based on p-values < 0.05. For both learning rates bs and bf, only the perturbed axis was added to the model with p = 2.98E-44 and p = 8.22E-14 respectively. This means that we did not observe a significant contribution of savings or anterograde effects on the learning rates.

Since the task uses multiple reach targets, one separate model could be fitted for each target unless one assumes full generalization between sequentially visited targets. To avoid this pitfall and corresponding increase in complexity, we fitted the model using epochs, motivated by the fact that in each epoch each target was visited one time. When we also tested the model for the case where full generalization happens between two consecutives targets, we found similar results as for the model trained on epochs (R2 between K and bs: 0.29, Supplementary Fig. 4).

We also validated the ability of our model to extract correlated learning parameters whenever present. For this, we simulated an additional set of data where the slow learning rate bs was imposed to be 0.3 times the steady-state Kalman gain (see Methods). Figure 6A shows that we were able to reconstruct such correlation with our model. Conversely, when no correlation between the slow learning rate and the Kalman gain was introduced in the dataset, the model does not introduce spurious correlation. Figure 6 overall confirms the validity of our model for the study of correlations between model parameters.

**Figure 6.**
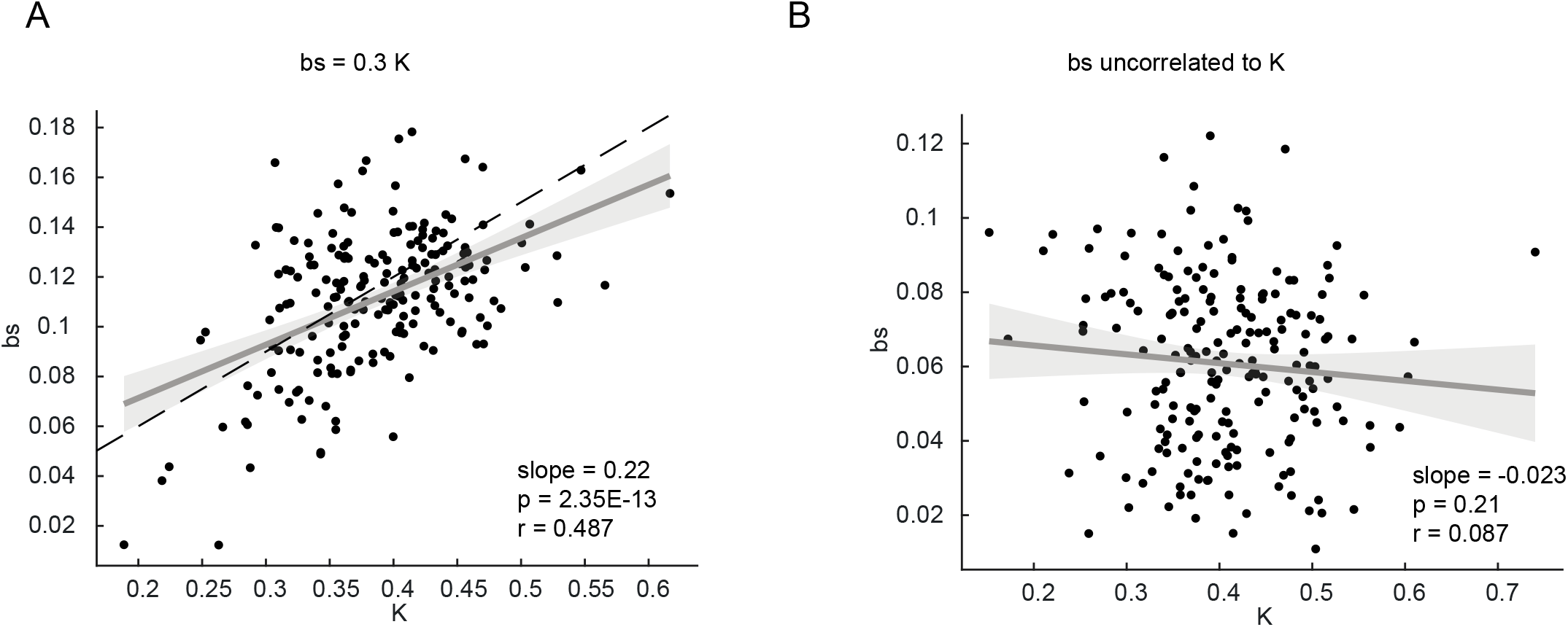
Recovery of simulated correlations between model parameters by the HMC algorithm. A) A correlation among bs and K was introduced in the surrogate data by setting bs equal to 0.3 times K (dashed black line). B) When no correlation between bs and K was added to the surrogate data, the HMC fitting algorithm did not introduce artificial spurious correlations.

## Discussion

We tested if observations and model assumptions about optimal integration (OI) theory for motor learning, as previously derived from one- or two-dimensional highly confined reach tasks, also hold true for largely unconstrained movements in three-dimensional space. We show that the rate of visuomotor adaptation under 3D stereoscopic vision depends on the plane in which a 2D visuomotor rotation perturbation is applied. Subjects adapted best to perturbations in the coronal plane followed by sagittal and horizontal planes. The rates of slow and fast learning correlated well with the Kalman gain when considering the variation across perturbation conditions, but not across subjects within a condition. We could separately assess the within and across condition parameters with our novel hierarchical Bayesian HMC fitting of the data. These results show that important aspects of OI theories of motor control derived from 2D movements generalize to the largely unconstrained 3D movements studied here, at least at the average population level, but do not extend to the individual subject level for any of the applied perturbations^20^.

### A novel two-state hierarchical HMC algorithm for precise estimation of motor learning parameters

We developed a novel HMC Bayesian fitting procedure with a multilevel hierarchical structure to take into account effects from multiple subjects performing a motor adaptation task in three different learning conditions, i.e., when different planes relative to the body were perturbed. Naturally, such a hierarchical approach shows lower goodness-of-fit to single-subject adaptation curves than algorithms that focus on single-subject data since the hyper-parameter at the population level (i.e. all the samples coming from a same distribution) act as regularization term. But multi-subject experiments are typically designed to determine the effects of experimental variations on the motor learning behavior independent of test subject. Hence, it is most important to best estimate the underlying learning parameters.

For the current experimental design, our procedure is better able to reconstruct actual model parameters than a state-of-the-art EM fitting algorithm^45^ or a non-hierarchical Bayesian HMC model, as revealed from the surrogate testing. This is due to the low identifiability of such two-state models for relatively short adaptation experiments comprising only a few dozen or even hundreds of trials per condition. The low identifiability is a result of random fluctuations in the shape of the adaptation curve and is easily associated with various learning parameters (see, for example, the outliers in Fig. 4). In the hierarchical model, each parameter is sampled from a population-level distribution, the parameters of which are also part of the model. This makes extreme values for the reconstructed parameter less likely.

Using our hierarchical model, we show that a two-state model fits the data significantly better than a single-state model. The two-state model is well established for subjects performing in a 2D environment^7,37^ and better explains learning mechanisms such as savings and anterograde interference in 2D settings^38^. As previously hypothesized^4,6^, the fast learning process can be partly associated to an explicit process and the slow learning process to an implicit process, both with anatomically distinct associated brain areas^6,19,43^. While it is plausible to assume that an interplay of explicit and implicit learning mechanisms is independent of the dimensionality of the task, to our knowledge, the two-state model to date has not been demonstrated for adaptation paradigms involving 3D movements. The fact that, in a 3D setting, we found a two-state model to better fit the adaptation data than a single-state model without introducing paradigms that explicitly enforce two-states dynamics (like re-learning after washout trials,^37^), might indicate that the explicit learning component in our three dimensional-paradigm is more pronounced than in classic two-dimensional paradigms. This finding is also in line with a recent study showing that VR visuomotor rotation tasks have a stronger explicit component than non-VR tasks ^8^. The results therefore further strengthen the association between fast and slow states with explicit and implicit mechanisms.

### Differences in estimated planning and measurement noises explain differences in learning rate

Motor learning studies are informative for motor rehabilitation protocols for patients that need to relearn or re-adapt existing motor schemes. These patients will necessarily perform their movements in a three-dimensional environment^55–58^. In our experiment, subjects adapted best to perturbations in the coronal plane, followed by sagittal and horizontal planes. Differences in the slow and fast learning rates between the planes were associated with different adaptation levels that were achieved during perturbation trials in our data. In the following, we will discuss how these anisotropies in adaptation are related to anisotropies in estimated planning and measurement noise, and, hence, could be explained by effects of gravity and depth perception, respectively.

The slowest adaptation in the horizontal plane is likely explained by an anisotropy in planning variability resulting from gravity. Previous studies suggested that movements along the vertical dimension have a higher level of planning variability associated with gravity compensation^33–36^. Optimally combining uncertainties on planning together with uncertainties on the estimation of the effector position here then means that sensory measurements should be given more credibility in vertical movements due to the higher planning variability. This leads to the prediction that adaptive corrections in the horizontal plane arise slower than corrections in the sagittal or coronal plane, since the latter contain the vertical dimension with the higher measurement credibility. Consistent with this view, perturbations in the horizontal plane in our data were associated with the slowest adaptation rates and planning variability was lowest in the horizontal plane, while measurement variability in the horizontal plane was close to the coronal plane (Fig5B).

The fastest adaptation rates in the coronal plane are likely explained by anisotropies in measurement variability resulting from visual depth perception. The coronal plane in our data yielded the lowest estimate of measurement noise and, correspondingly, subjects adapted faster to perturbations in this plane (Fig. 5B). In our model, measurement noise represents the reliability of the subject measurements of the cursor position, i.e., the visual feedback about the movement, since the cursor determines the motor goal. Subjects have to rely mainly on stereoscopy to estimate visual depth and visual localization is less precise in depth^14,32^. Since depth is not perturbed by a rotation in the coronal plane, measurement error is least in the coronal condition and adaptation should be fastest. On the other hand, visuomotor rotations in the sagittal and horizontal planes both resulted in perturbations of cursor depth compared to baseline. Corresponding to this, other studies also showed that subjects tend to adapt worse when the measurement variability (or sensory nose) was increased by practically blurring the error feedback^11,13,21^.

### Optimal integration theory in 3D

Previous studies^11,20,21,26,31^, provided evidence for the OI theory during visuomotor adaptation. They showed that the planning component of motor noise correlates positively with the adaptation rate while the measurement component correlates negatively, at least when the respective noise component is experimentally kept constant. While the adaptation data from our 3D movements yielded similar trends, the correlation of slow learning rate with either measurement- or planning-variability was weak. For an optimal learner, the Kalman gain is indicative of the level of correction attributed to the experienced error 11,21,31. This Kalman gain positively correlates with the slow learning rate in our data, supporting the idea that subjects optimally integrate the information relative to the internal state with experienced (visual) feedback. However, and as opposed to what was suggested in previous studies^20,31^, we observe the correlation between Kalman gain and learning rates only across different perturbation conditions. Within each perturbation condition and across individual subjects, there was no relationship between Kalman gain and learning rates.

## Conclusion

Here, we develop a novel HMC fitting procedure to correlate learning rates and noise levels estimated with the same model and without the need to experimentally manipulating these error variabilities. By applying this model to a sensorimotor adaptation task with naturalistic movements in 3D, we found that the rate of adaptation with which subjects counteract applied visuomotor rotations, depends on the statistics of the combined errors with which subjects plan and perceive movements. These errors differ between different axes relative to the body, likely due to gravity compensation during planning- and perception-induced anisotropies. This insight could be used in motor rehabilitation strategies by specifically targeting the orientation of the body relative to the target while performing reaching movements to accelerate learning.

## Acknowledgments

German Research Foundation (DFG, Germany, grant number FOR-1847-GA1475-B2), received by AG. German Research Foundation (DFG, Germany, grant number SFB-889), received by AG. Federal Ministry for Education and Research (BMBF, Germany, grant number 01GQ1005C), received by AG. Federal Ministry for Education and Research (BMBF, Germany, grant numbers 01GQ0814), received by AG.

## Supplementary Figures

**Supplementary Figure 1.**
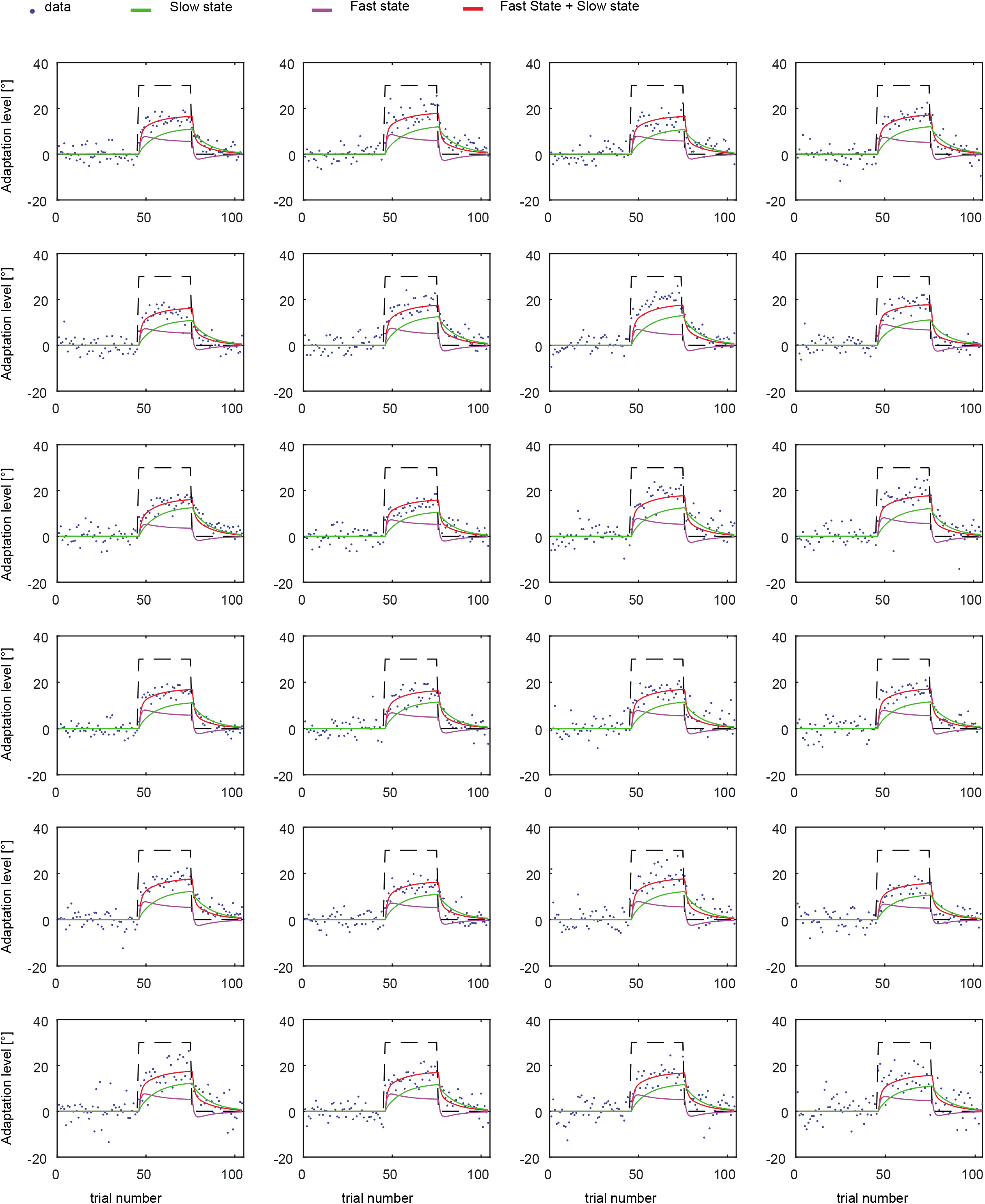
Results of fitting the hierarchical HMC model to the adaptation profile of each subject (n = 24) with perturbation applied to the Sagittal plane.

**Supplementary Figure 2.**
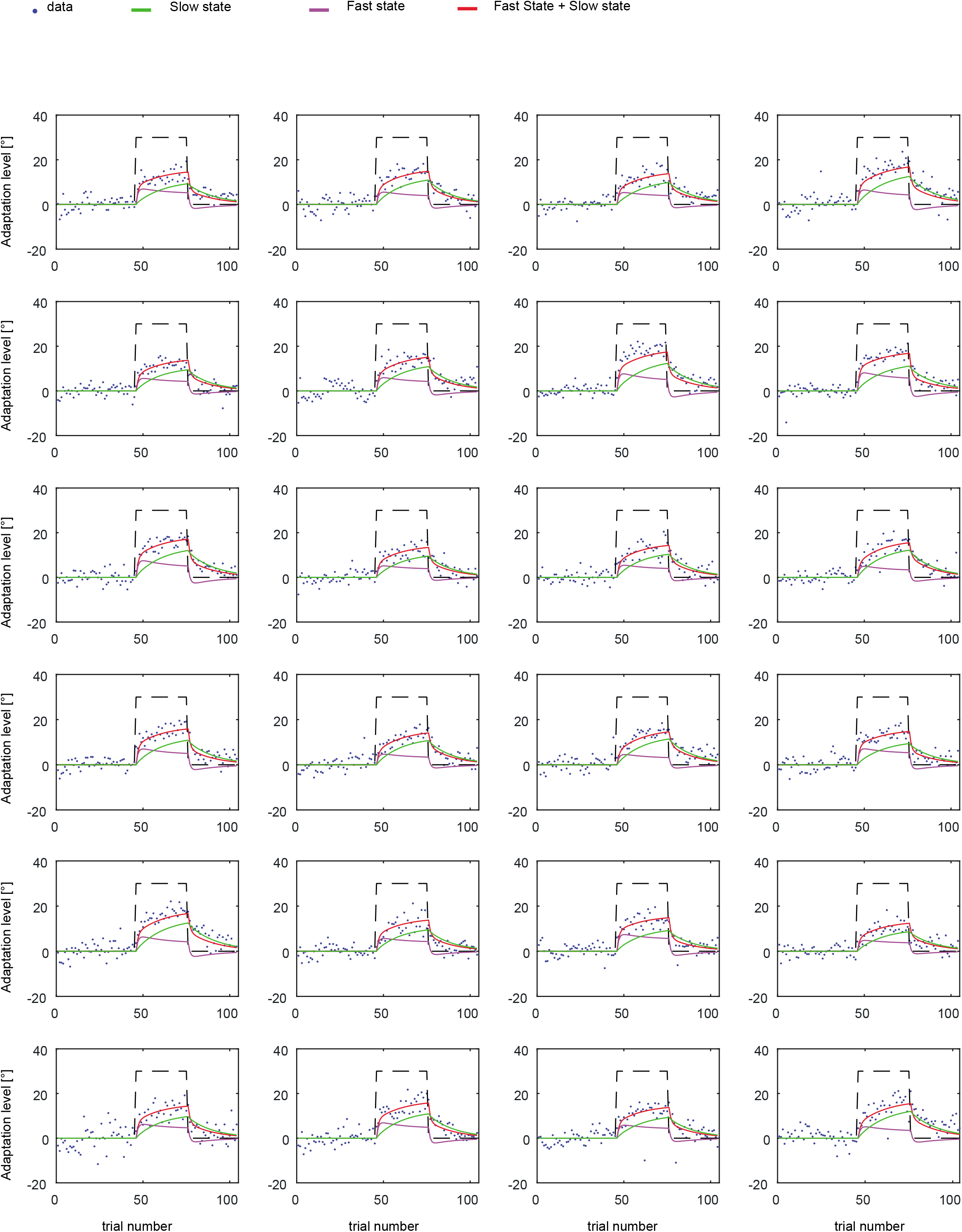
Results of fitting the hierarchical HMC model to the adaptation profile of each subject (n = 24) with perturbation applied to the horizontal plane.

**Supplementary Figure 3.**
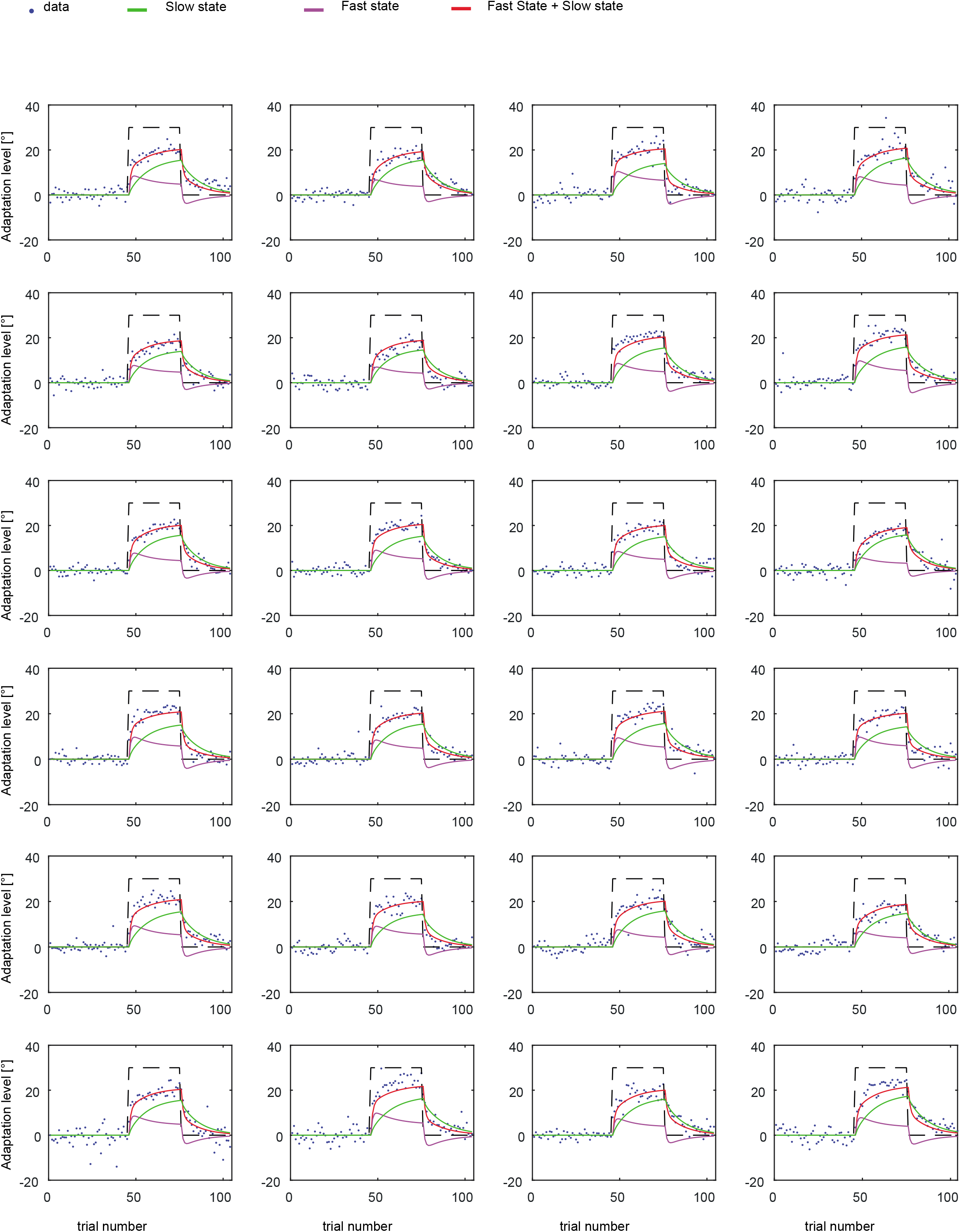
Results of fitting the hierarchical HMC model to the adaptation profile of each subject (n = 24) with perturbation applied to the coronal plane.

**Supplementary Figure 4.**
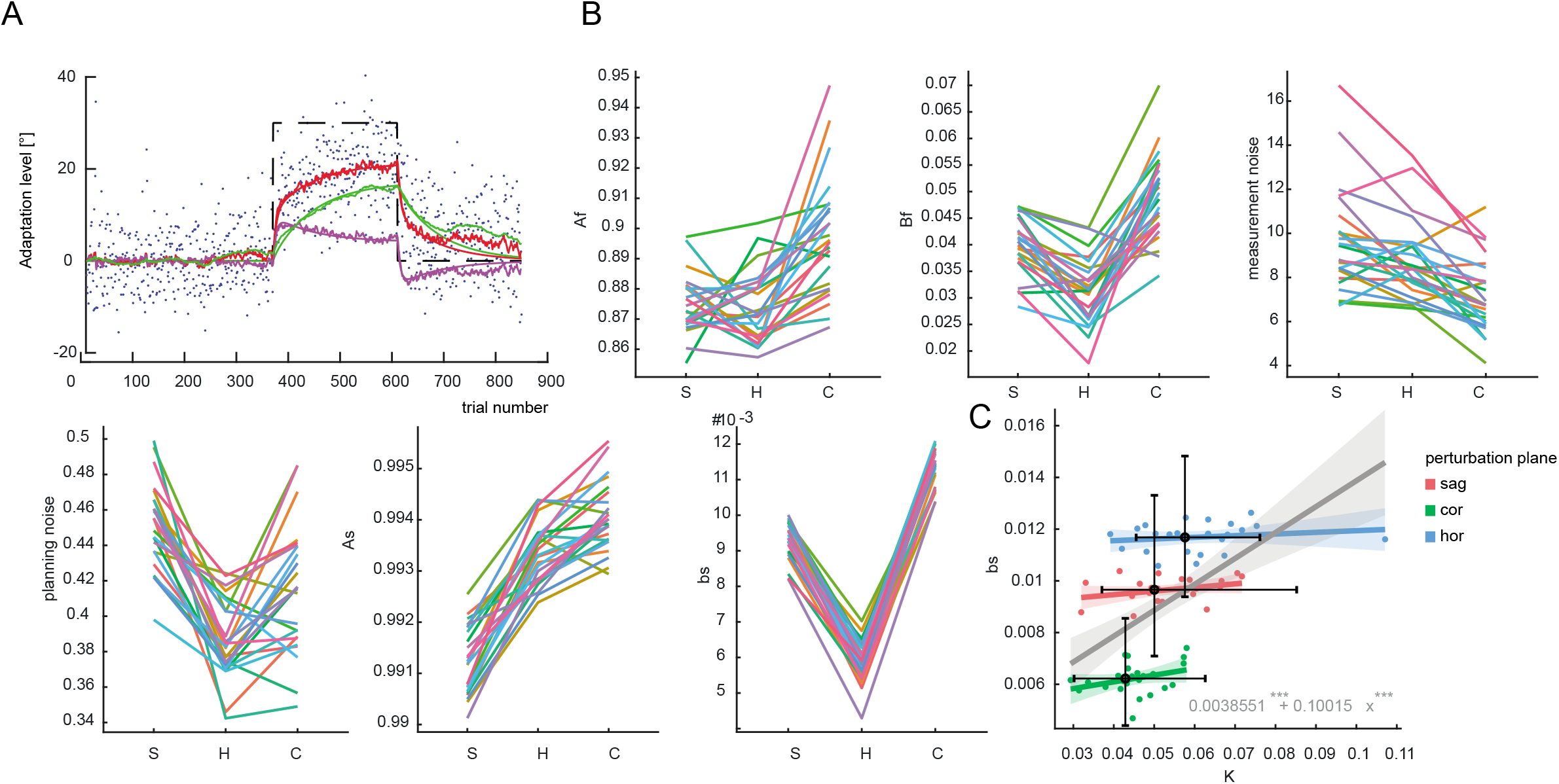
Similar to figure 5 but with the model applied on a trial base rather than epoch base (no average over 8 consecutive trials). A) Example of single subject time course of adaptation with 2-state model fits. B) Extrapolated model parameter for all the subjects (different colors). C) Slow learning rate (bs) vs Kalman gain relationship.

